# Simultaneous recordings link intracranial and scalp correlates of somatosensory awareness in humans

**DOI:** 10.64898/2026.10.03.756393

**Authors:** Ezequiel Mikulan, Flavia Maria Zauli, Simone Russo, Francesca Mannini, Maria Del Vecchio, Davide Albertini, Piergiorgio d’Orio, Lino Nobili, Ivana Sartori, Marcello Massimini, Pietro Avanzini, Andrea Pigorini

**Affiliations:** Department of Biomedical and Clinical Sciences “L. Sacco”. University of Milan, Milan, Italy; Department of Health Sciences. University of Milan, Milan, Italy; ASST GOM Niguarda. Piazza dell’Ospedale Maggiore 3, Milan, Italy; Theoretical and Applied Neuroscience PhD program, University of Camerino; Wallace H Coulter Department of Biomedical Engineering, Georgia Institute of Technology and Emory University, Atlanta, GA, USA; Institute of Neuroscience, Consiglio Nazionale delle Ricerche, Parma, Italy; Istituto G. Gaslini. Genoa, Italy; Department of Neuroscience, DINOGMI, University of Genoa, Genoa, Italy; Istituto Di Ricovero e Cura a Carattere Scientifico, Fondazione Don Carlo Gnocchi ONLUS, Milan, Italy; Department of Biomedical, Surgical and Dental Sciences. University of Milan, Milan, Italy; UOC Maxillo-facial Surgery and dentistry, Fondazione IRCCS Cà Granda, Ospedale Maggiore Policlinico, Milan, Italy

**Keywords:** Awareness, Perception, EEG, SEEG, Simultaneous Recordings, Somatosensory

## Abstract

The electrophysiological events underlying the conscious perception of touch in humans remain elusive. Scalp-EEG studies have identified a negative deflection peaking 120-200 ms after stimulus onset, known as the N140. This component, recently reinterpreted as the somatosensory awareness negativity (N140/SAN), has been proposed as a putative marker of tactile awareness, but its underlying neuronal dynamics are still unknown. Intracranial studies have shown that responses to median nerve stimulation include, after an initial phasic activation mainly localized to primary somatosensory cortices, a predominantly insulo-opercular tonic broadband high-frequency activity (BHA; 50-150 Hz) that has been linked to tactile awareness. Yet these scalp and intracranial signatures have never been directly linked. Using simultaneous high-density scalp EEG and intracranial EEG recordings, we show that the N140/SAN is the scalp manifestation of the insulo-opercular local field potential that follows the BHA response to stimulation. By bridging invasive and non-invasive recordings our results identify the neural origins of the N140/SAN and connect scalp and intracranial correlates of somatosensory awareness.

**Significance statement:** Understanding the neural correlates of somatosensory perception remains a central challenge in neuroscience. This study bridges macroscale and mesoscale perspectives by linking the scalp N140/SAN—a putative marker of somatosensory awareness—to intracranial local field potentials (LFPs) and broadband high-frequency activity (BHA, 50-150 Hz) in the parietal operculum and posterior insula. These findings reveal the generators of the N140/SAN and link scalp and intracranial accounts of somatosensory awareness.

## Introduction

The chain of neuronal events underlying somatosensation have been investigated since the early days of modern neuroscience^1–5^. Following transduction by tactile peripheral receptors, neuronal activity is transmitted through distinct sensory nerve fibers, ascends passing through thalamic relays, and activates the cerebral cortex via primary somatosensory areas (SI: Brodmann’s areas 3a, 3b, 1 and 2; postcentral gyrus). From there, somatosensory information travels to higher-order centers in the cortex, including the posterior parietal cortex and the secondary somatosensory cortex (SII: Brodmann’s areas 40 and 43, in the parietal operculum), where more integrated processing takes place^6^. Within this complex chain of events, when and where a delivered tactile stimulus turns into a consciously perceived one is still unclear^7^.

Growing evidence from electroencephalography (EEG) and magnetoencephalography (MEG) studies has focused on a negative deflection observed on scalp EEG/MEG event-related potentials (ERPs), the long-known N140 component^8^, which peaks ∼140 ms after stimulus onset and has been recently reframed as the Somatosensory Awareness Negativity^7,9^ (here referred to as N140/SAN; see below). This component (i) emerges ∼80 ms after stimulus delivery; (ii) is typically bilateral over fronto-temporal electrodes, but tends to be larger contralaterally to the stimulation side; (iii) is present for both target and non-target stimuli in active and passive paradigms; and (iv) has been associated with stimulus perception^9,10^. EEG/MEG studies have estimated the sources of these activations in SII, the posterior parietal cortex, and the paracentral lobule^11–16^.

At the mesoscale, stereoelectroencephalography (SEEG) shows that somatosensory stimulation produces, after an initial phasic activation mostly located in primary somatosensory cortices (linked to stimulus encoding), a long-lasting increase in broadband high-frequency activity (BHA; 50-150 Hz), termed the tonic response. This activation (i) peaks beyond ∼60 ms and lasts up to ∼250 ms, (ii) is mostly located in the parietal operculum and posterior insular cortex^17,18^, (iii) is bilateral and (iv) does not follow a somatotopic organization^19,20^. Importantly, it has been shown that lesions to these areas can produce deficits of tactile awareness^18^ and a recent study has shown that tonic BHA responses correlate with stimulus awareness in near-sensory-threshold tasks^21^. Notably, BHA, also known as high-gamma or epsilon band, has been associated with neuronal spiking and dendritic processes and is often used as a proxy for the former^22,23^.

The scalp N140/SAN and the intracranial insulo-opercular BHA response show similar time courses and seem to originate from overlapping anatomical regions, but they have never been linked. Here, we hypothesized a relationship between these two putative markers of somatosensory perception and directly tested it with simultaneous recordings of high-density EEG (HD-EEG) and SEEG, leveraging, concurrently, the whole-brain coverage of the former and the high spatiotemporal resolution and signal-to-noise-ratio (SNR) of the latter.

Our results demonstrate that the N140/SAN is the scalp manifestation of the intracranial local field potential (LFP) that follows the insulo-opercular BHA response to somatosensory stimulation. These results connect two putative markers of somatosensory perception and their respective bodies of literature, and pave the way for a multiscale understanding of somatosensory perception in humans.

## Results

Simultaneous recordings of EEG (256 channels) and SEEG during median nerve electrical stimulation were obtained from 8 subjects (Figure 1.a-c; Supplementary Table 1). All subjects were stimulated on the wrist contralateral to the intracranial implant and 2 subjects were also stimulated on the ipsilateral side (S2 and S4), constituting a total of 10 sessions. From each session we extracted the time-locked EEG, LFP and BHA (50-150 Hz) responses (Figure 1.d). Out of the 1306 SEEG recording contacts, 27 showed tonic BHA responses as previously defined^17^ (Figure 1.c; 2.06%, average number per subject = 3.37 ± 2.06, range = 1-7). In subjects with more than one BHA responsive contact, these were often immediate neighbours 1.5 mm apart (Figure 1.e-f). We therefore selected the site with the highest response magnitude per subject to avoid including partially redundant information, and to retain a single contact for group level analyses (Figure 1.f-g). This avoided introducing a nested data structure that would have required explicit modeling of within-subject dependence^24^ (see Methods).

**Figure 1.**
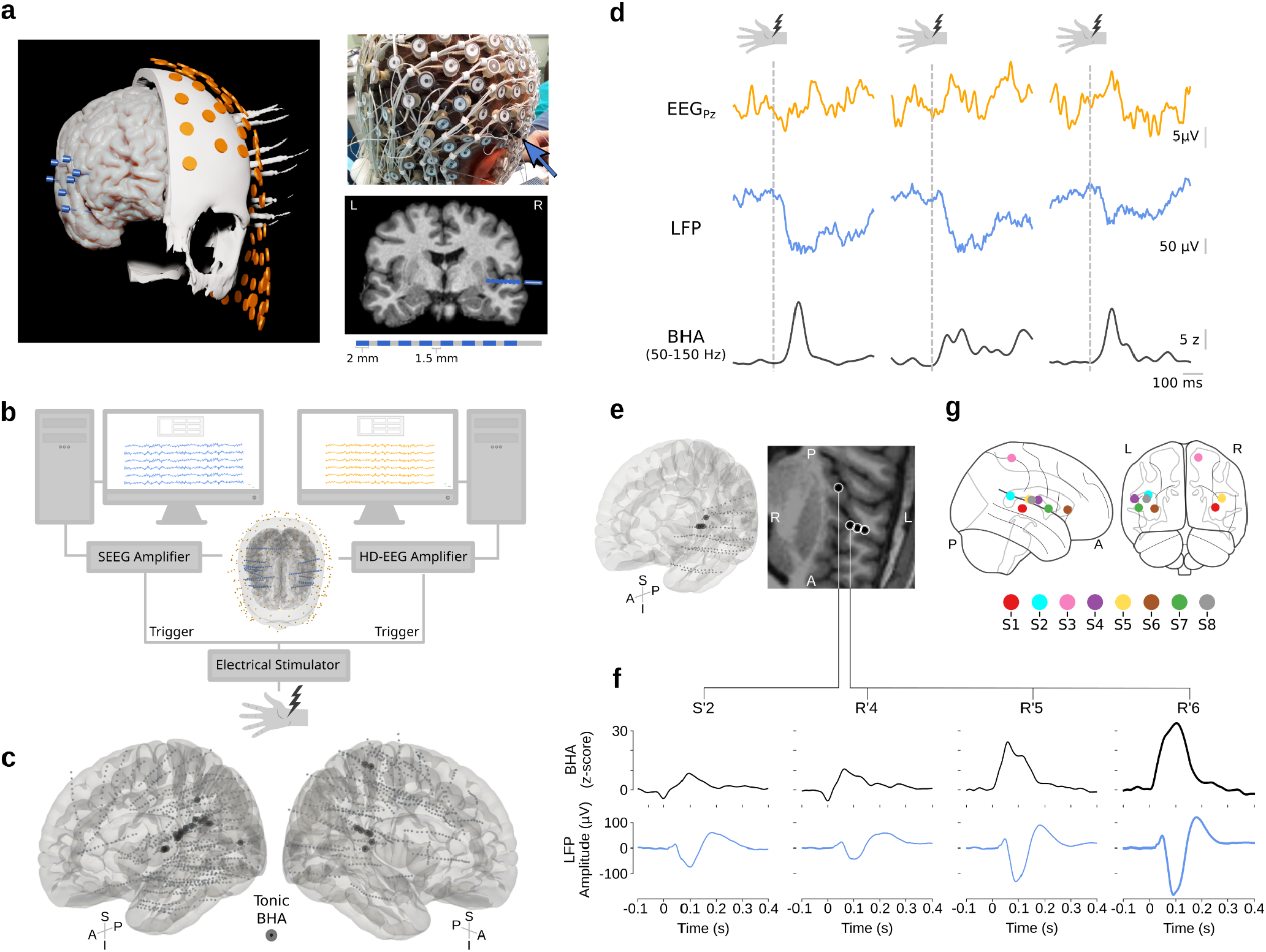
Simultaneous SEEG and HD-EEG recordings in humans. **a**) Coregistration. *Left*: 3D reconstruction of the brain, skull, EEG (orange) and SEEG (blue) recording electrodes of a subject undergoing simultaneous SEEG and HD-EEG. *Top right*: Photograph showing the HD-EEG electrodes and the external segment of the SEEG implant (arrow). *Bottom right*: T1-weighted MRI slice and recording contacts (blue dots) of an SEEG electrode (top) and its schematic representation (bottom). **b**) Experimental setup. An electrical stimulator delivered pulses to the median nerve while sending triggers to the two amplifiers for offline synchronization. **c)** 3D representation of the normalized positions (MNI152) of all recorded channels (grey points) and contacts showing sustained BHA (black). A larger version of this image is provided in Supplementary Figure 1. **d)** Single trial responses as observed from the scalp (orange), intracranially (blue) and the corresponding BHA extracted from the latter (black). **e)** Contacts showing sustained BHA from a single-subject on a 3D surface (left) and MRI slice (a maximum intensity projection of one slice was applied in order to show R’4 together with the other three channels). **f)** Average BHA (black) and LFP (blue) for all channels showing sustained BHA for a representative subject (same as panel e). The channel selected for subsequent analyses is highlighted (R’6). **g)** Normalized positions of the selected channels for each subject.

Average EEG responses showed the SAN/N140 component which peaked at 125 ± 35 ms with the typical bilateral negativity larger contralateral to the stimulation site (Figure 2.a-b). SEEG tonic BHA responses peaked at 96 ± 19 ms and were followed by a deflection of the LFP that peaked at 124 ± 28 ms (Figure 2.c). As illustrated in Figure 2d, the EEG response in a region of interest (ROI) encompassing the SAN/N140 scalp territory contralateral to the stimulation site was closely aligned in time with the LFP.

**Figure 2.**
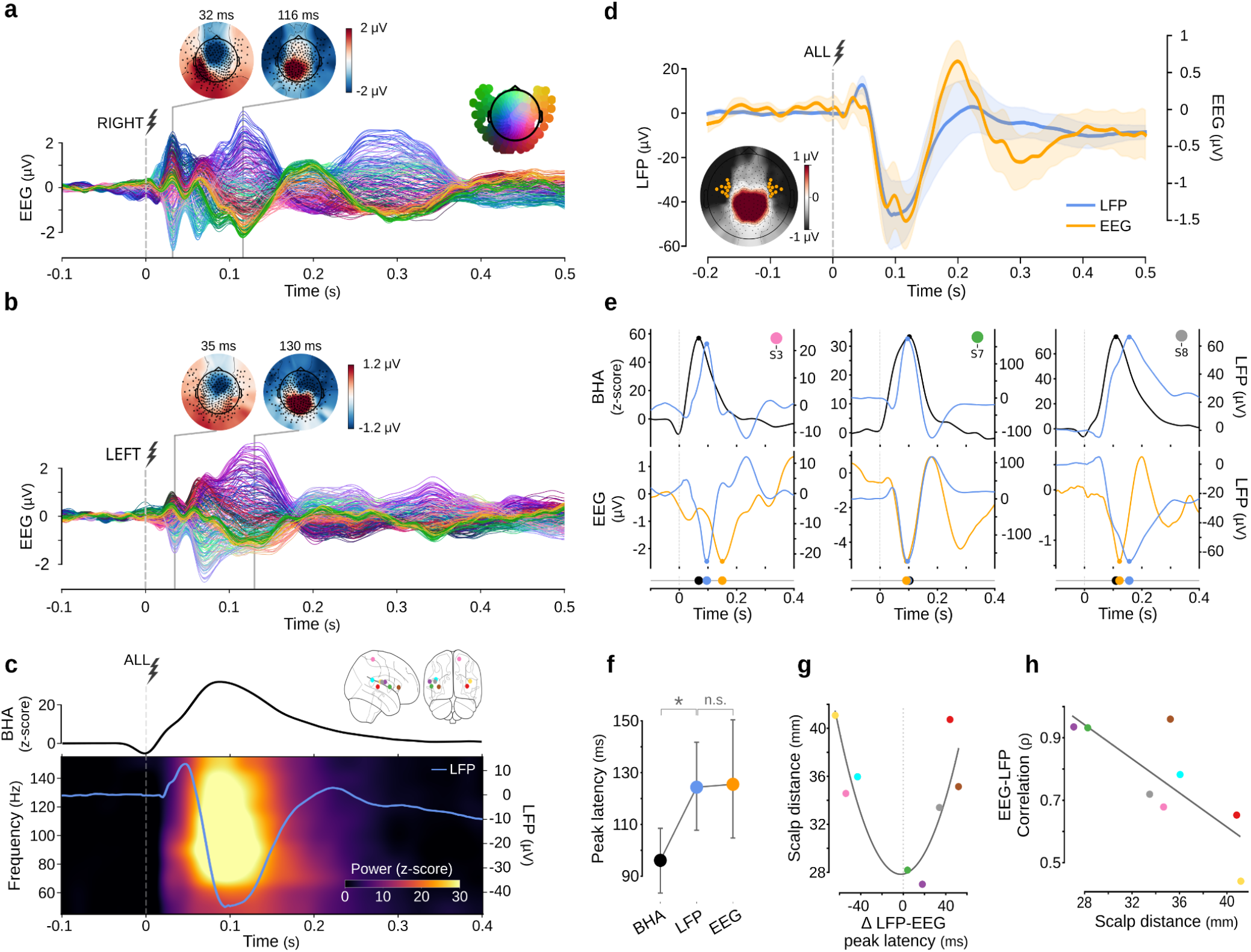
Simultaneous SEEG and HD-EEG recordings link intracranial BHA and LFP activity with the scalp N140/SAN. **a**) Average scalp EEG somatosensory evoked potentials across subjects when stimulating the right median nerve. Colors indicate channel positions as displayed in the inset diagram. Topographies indicate voltage distributions on the scalp at the early and late signal peaks within the temporal window of interest. **b)** Same as panel d, for the stimulation of the left median nerve. **c)** Average intracranial responses to median nerve stimulation, including both left and right stimulations. Top: time course of BHA (50-150 Hz). Bottom: time-frequency representation (color gradient) and average LFP responses (blue trace). **d)** Average EEG and LFP responses. EEG traces correspond to the average of the channels belonging to the left or right ROI, selected contralaterally to the stimulation site by session. Shaded areas represent the standard error of the mean. Inset topography shows average voltage distribution in the interval from 70 to 150 ms (FWHM) and EEG channels of the left and right ROIs. **e)** Representative single-subject (columns) average BHA (black), LFP (blue) and EEG-ROI (orange) responses. Traces are shown with a 30 ms moving average smoothing. Dots on top of the traces indicate peak latencies and are summarized on the timeline at the bottom. **f)** Average peak latency (circles) and 95% confidence interval (lines) for BHA, LFP and EEG responses across sessions. The asterisk indicates *p* < 0.05 and n.s. *p* > 0.05. **g)** EEG-LFP difference in peak latency and distance to the skin from the corresponding intracranial contact by subject with quadratic regression curve (line). **h)** Maximum of the cross-correlation between LFP-EEG (ρ) and distance from the scalp of each intracranial contact by subject. Colors in panels c (inset), g and h represent subjects as depicted in Figure1.g.

We first investigated the temporal relationships comparing peak timings between BHA, LFP and EEG (see Figure 2.e for three representative subjects and the Supplementary Figure 2 for all single-subject plots). This analysis showed that BHA responses preceded the LFP by ∼28 ms (t_(7)_ = −3.19, *p_corr_*= 0.03; paired t-test) whereas the ∼ 1 ms difference between LFP and EEG was not statistically significant (Figure 2.f; t_(7)_ = −0.07, *p_corr_* = 1, paired t-test; Supplementary Table 2).

Although peak latencies between LFP and EEG were comparable and their difference was centered around 0, within-subject LFP-EEG latency differences (Δ LFP-EEG peak latency) showed variability (2 subjects within ± 20 ms, 3 subjects > 20 ms, 3 subjects < −20 ms; Figure 2.g). We therefore tested whether this variability was related to contact depth by computing the minimum Euclidean distance between the contact’s location, in the subject’s native anatomical MRI space, and the 3D reconstruction of the subject’s scalp. Since differences could be either positive or negative, we used a quadratic regression analysis (*f*(*x*) = *a* + *b* · *x* + *c* · *x*^2^), and verified that the results held in a linear regression on the absolute Δ LFP-EEG peak latency. This analysis showed that larger timing differences corresponded to deeper contacts (Figure 2.g; *a* = 27.84, *p* = 1.12e^6^, *b* = 15.83, *p* = 0.58, *c* = 3558.01, *p*= 0.006; R²_adj_ = 0.63; n = 8; *vertex* = [-0.002 s, 27.82 mm]; Supplementary Table 3).

Likewise, we tested whether waveform similarity between LFP and EEG traces was also related to the distance to the scalp, by computing the maximum of the LFP-EEG cross-correlation (to allow for within-subject timing differences). The average correlation (ρ) was 0.76 ± 0.17 and the average lag was −3 ± 41 ms (Supplementary Table 4). Linear regression indicated that LFP-EEG similarity decreased with distance to scalp (Figure 2.h; β = −0.27 per cm, CI_95%_ [−0.49, −0.055], *p* = 0.02; R²_adj_ = 0.54; n = 8; Supplementary Table 5).

Together, these findings indicate that BHA responses precede the LFP, while LFP and EEG show comparable time courses, with high alignment and similarity that depend on spatial proximity.

To assess the spatial correspondence between the scalp N140/SAN and the tonic intracranial BHA responses we leveraged the high spatial sampling of our scalp recordings to perform source reconstruction. Spatial variability in source activity (source ***σ***), a metric analogous to the sensor-level global mean field power (GMFP) ^25,26^, showed three discrete peaks (which were also evident in the unprojected EEG averages, see Figure 2.a-b), with high spatial correlation among themselves (Figure 3.a), suggesting the successive reactivation of a similar spatial pattern (Figure 3.b). To test the similarity between the whole-brain map of the projected EEG N140/SAN activity and a whole-brain map of intracranial BHA, we compared our data with the dataset from a previous study, including 99 implanted epileptic patients^17^. Specifically, from our hd-EEG data we extracted the top 15 % of reconstructed sources in the window from 70 to 150 ms (FWHM of the EEG response in the ROI of the N140/SAN; ROI FWHM; see Figure 2.d) and compared it to a binarized density map of intracranial contacts showing sustained BHA (Figure 3.c). The similarity between these two maps was statistically significant (Dice Similarity = 0.34, *p* = 0.03; number of permutations = 5000; spatial autocorrelation-preserving null model; Supplementary Figure 3; see also the Supplementary Notebooks in the Supplementary Materials for an interactive visualization of source modeling results).

**Figure 3.**
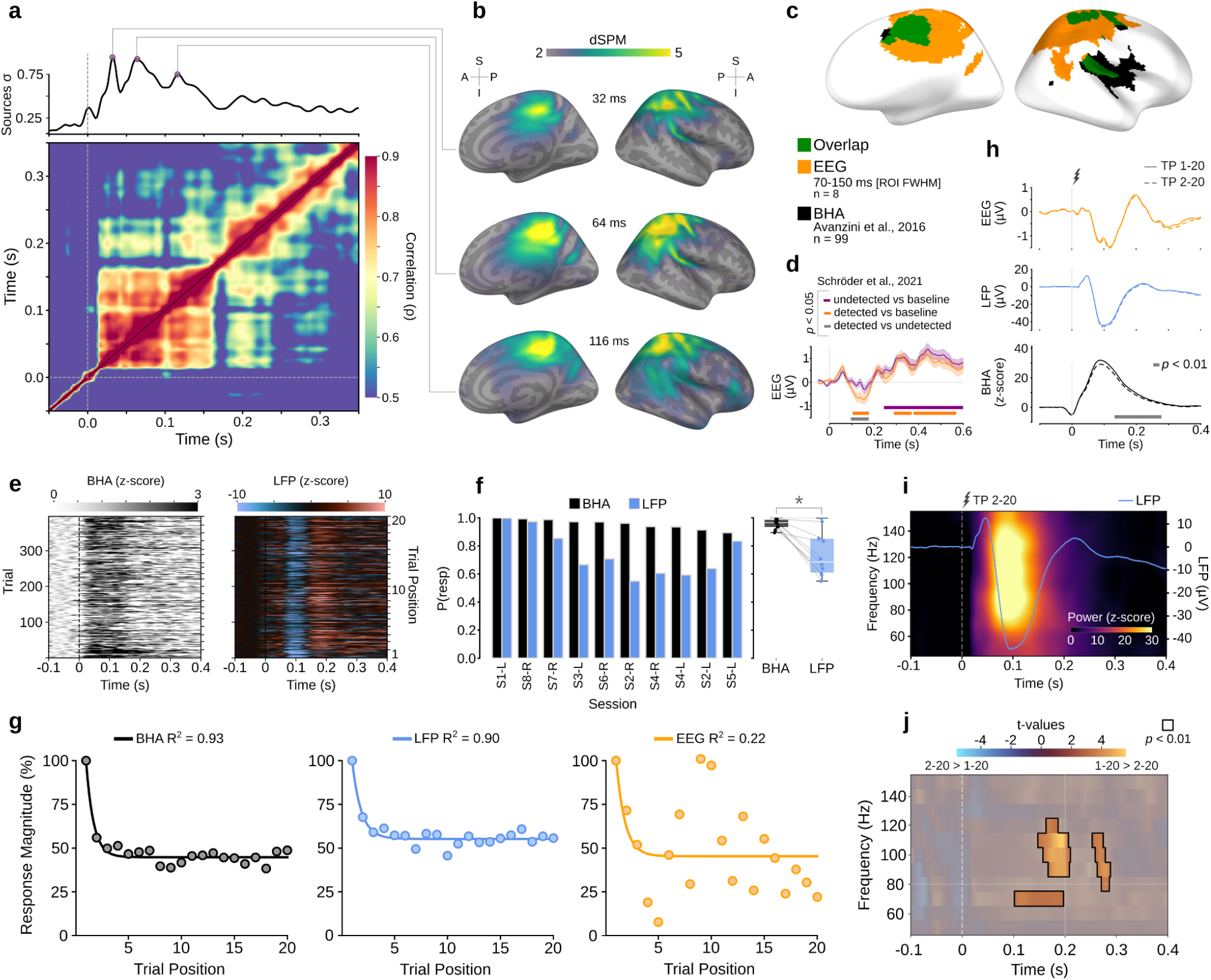
Source modelling, near-threshold analysis, single-trial response proportions and saliency. **a**) *Top:* Standard deviation across vertices from the source localization of the average scalp response. Dots indicate peaks. *Bottom:* correlation matrix of average source activity across timepoints. **b)** Surface representation of source activity at the peak timepoints from panel a (± 10 ms). **c)** Top 15 % of EEG source localized activations (orange) in the interval from 70 to 150 ms (FWHM of the EEG-ROI response), contralateral to the stimulation side; topography of sustained BHA (50-150 Hz) responses (black) recreated from Avanzini et al, 2016; and corresponding overlap (green). Source activity in panels b and c was mapped to a single hemisphere by left-right morphing. Dice Similarity between maps was statistically significant (*p* < 0.05; spatial autocorrelation-preserving null model). **d)** Evoked responses to near-threshold detected (orange) and undetected (purple) stimuli in a contrastive task (data from Schröder et al., 2021). Colored segments indicate *p* < 0.05 when testing either between conditions or against baseline. **e)** Single trial BHA (left) and LFP (right) responses from a representative subject with trials ordered by position in the stimulation block. **f)** Proportion of trials showing responses greater than 2σ_baseline_, by subject (left) and group (right). **g)** Average response magnitude in the interval 70-150 ms (FWHM of the EEG-ROI response) as a function of trial position in the stimulation block for BHA (left), LFP (center) and EEG (right) with respective fitted exponential decay functions. **h)** Comparison of evoked responses for EEG (top), LFP (middle), and BHA (bottom) when including (TP 1-20) or excluding (TP 2-20) the first trial of each stimulation block. The colored segment indicates *p* < 0.05. **i)** Average time-frequency representation of SEEG responses for TP 2-20. See Figure 2.c for the corresponding plot for TP 1-20. **j)** Comparison of time-frequency representations between TP 1-20 and TP 2-20. Reference lines are shown at 80 Hz and 200 ms. Colors represent t-values and highlighted non-greyed out segments indicate *p* < 0.01.

Since our experiment used a supra-threshold, no-report paradigm, it did not allow to contrast between *detected* and *undetected* stimuli at the verge of sensory threshold, which has been done in studies targeting the SAN^7,9,27^. Therefore, to relate our findings to this literature we reanalyzed data from an independent study^9^. Specifically, we compared the N140/SAN responses to *detected* and *undetected* near-threshold stimuli (Figure 3.d) by testing the differences between conditions and each one against baseline. Permutation cluster tests showed that, in the temporal window of the N140/SAN, responses were different between *detected* and *undetected* stimuli (cluster from 95 to 175 ms, *p* = 0.017) but only the *detected* condition was significantly different from baseline activity (cluster from 103 to 177 ms, *p* = 0.043). Both conditions differed significantly from baseline in a later window, but not from each other in this interval (∼300 to 600 ms; see Supplementary Materials).

Having established the link between the SAN/N140 and intracranial responses, we leveraged the high SNR of invasive recordings to address open questions about the scalp component that are otherwise difficult to tackle. A fundamental open question is whether the N140/SAN correlates with stimulus perception at the single trial level or only on the average^7^. To lay the groundwork for answering this question, we assessed the detectability of intracranial responses at the single-trial level. We found that, with our supra-threshold experimental design, trials exceeding 2*σ_baseline_* amplitude were present in 95 ± 3 % of trials across subjects in BHA signals, and in 74 ± 16 % in the LFP, a difference that was statistically significant (Figure 3.e-f; *W* = 0; *p* = 0.0019; median difference = 0.269; Wilcoxon signed-rank test; Supplementary Table 6).

Another fundamental open question is how stimulus saliency affects the characteristics of the observed responses^7^. Therefore, we characterized the saliency/habituation effects^28^ (i.e. an increased response magnitude due to salient stimuli or a decrease due to repeated stimulation). In line with previous EEG literature^29^, we compared an exponential decay function (*f*(*x*) = *a* + *b* · *exp*(− *c* · *x*)) and a hyperbolic decay function (*f*(*x*) = *a* + *b*/*x*), fitted to the average response magnitude in the interval 70-150 ms (ROI FWHM) as a function of trial position (TP) in the stimulation block (Figure 3.g; Supplementary Table 7; Supplementary Figure 4). Results showed that BHA responses were better captured by the exponential decay, quickly approaching a plateau of ∼45 % the initial magnitude (*a* = 44.81 ± 0.89, *p* < 1e^-16^; *b* = 228.04 ± 64.81, *p* = 0.002; *c* = 1.42 ± 0.26, *p* = 5.213e^-5^; R^2^=0.93), with strong support over the hyperbolic decay (ΔAICc = 10.15). For LFP, the exponential decay also provided a good fit, approaching ∼ 55% of the initial magnitude but with a slower rate than BHA (*a* = 55.26 ± 0.90, *p* = 1e^-16^; *b* = 144.47 ± 39.47, *p* = 0.001; *c* = 1.17 ± 0.24, *p* = 0.0001; R^2^=0.90), again with strong support over the hyperbolic decay (ΔAICc = 6.45). Conversely, EEG responses were not well captured by either decay function (exponential: *a* = 44.23 ± 6.25, *p* = 1.88e^-6^; *b* = 238.49 ± 477.92, *p* = 0.62; *c* = 1.45 ± 1.88, *p* = 0.45; R^2^=0.21, ΔAICc = 1.51), arguably due to its lower SNR. Notably, the larger response magnitude of trials in first position was already evident at the single-subject level for both BHA and LFP (Figure 3.e).

Given the observed saliency effect, we repeated all previous analyses but excluding the first trial of each stimulation block. Although exact values differed slightly, results were unchanged (see Supplementary Analysis 1).

Finally, to assess whether and how saliency can shape the observed responses, we tested the effect of including the salient first-position trial of each block (TP 1–20), as opposed to excluding it (TP 2–20), on the averaged time courses. EEG and LFP showed comparable waveforms, while BHA responses were significantly larger in TP 1-20 (Figure 3.h), as evidenced by a cluster ranging from 132 to 278 ms (*p* = 0.0039; one-sample permutation cluster test on paired differences). To further characterize this effect, we compared the time-frequency representations and found that responses in TP 2-20 were temporally and spectrally more confined (Figure 3.i-j; see also Figure 2.c and Supplementary Figure 5 for a side-by-side representation), as evidenced by three clusters with higher power values in TP 1-20 (from 102 to 196 ms, 70 Hz, *p* = 0.007; from 148 to 210 ms, 90 to 120 Hz, *p* = 0.003; and from 254 to 288 ms, 80 to 110 Hz, *p* = 0.007; permutation cluster test on paired differences).

The data and code to reproduce the presented results are publicly available (see Methods and Supplementary Materials).

## Discussion

The quest to identify the intracranial generators of scalp event-related potentials (ERPs) has been traditionally performed using either source modeling techniques or comparisons with asynchronous intracranial recordings^30–32^, with the latter regarded as the gold standard^33^. The indirect nature of these approaches limits the precision with which the relationship between scalp potentials and their intracranial counterparts can be assessed. The present study leverages scalp and intracerebral signals acquired simultaneously to compare them not only in space, but also in time, with millisecond resolution^34^.

Converging studies on the neural correlates of consciousness (NCCs) have pointed to a putative marker, the perceptual awareness negativity (PAN), whose modality-specific components share common features, including latency, polarity and scalp location above the respective sensory cortices^7^. In the somatosensory domain, the SAN (i.e. the somatosensory variant of the PAN) is a reinterpretation^7^ of the long-known N140 component^8^ of the ERP response evoked by tactile stimulation. Are they different? The N140 was first reported in 1977 by Desmedt and Robertson^8^ and has since been a physiological and psychological object of study in over a hundred papers (see Supplementary Analysis 2). In the more recent search for the correlates of consciousness the N140 was redefined as the SAN^7^, but has, however, been most specifically associated with contrastive studies using active tasks and near-threshold stimulations^7,9,27^ (see below). Still, the definition of the neural event of interest should not be conflated with its operationalization. Notably, the waveforms, latencies, and topographies observed in contrastive studies when stimuli are perceived closely resemble those observed in non-contrastive paradigms^7,9,27^, as also evident in our results (Figure 2.a-b), whereas activity in the same time window remains near baseline when stimuli are not perceived (Figure 3.d). To our knowledge, this issue has not been addressed through a direct comparison. Consequently, whether the N140 and the SAN represent two different phenomena or are two views of the same neural event remains to be elucidated, and will be treated jointly below.

More generally, despite decades of research, the neuronal dynamics underlying these components are still insufficiently characterized, with evidence largely limited to indirect localizatory descriptions. The origins of the long-latency components of SEPs have been previously investigated by means of EEG/MEG source localization. Multiple studies found that scalp signals between 80 and 180 ms after somatosensory stimulation originate from SII, the posterior parietal cortex, and the mesial cortex^11,13–16,27,35^. However, because these methods infer neural sources from scalp-level recordings, they do not directly capture the relationship between scalp and intracranial dynamics, and their detection of activity above ∼30 Hz is heavily limited by noise^36–38^.

To the best of our knowledge, the only previous attempt to assess the within-subject direct relationship between intracranial and scalp recordings of SEPs was carried out by Allison and colleagues in 1992^39^. The authors compared scalp-only, intracranial-only and low-density simultaneous recordings and speculated that the N120-P120 scalp component recorded from the scalp might correspond to the intracranial perisylvian N100-P100 LFP component; see also^40^. They also proposed that the scalp N140 was likely generated bilaterally in the frontal lobes—specifically orbitofrontal, mesial, and lateral regions. Additionally, they found large potentials in the mesial central cortex at 130 ms which they considered possible contributors to the scalp N140. However, this study only assessed LFP components and results were not conclusive, as most of the simultaneous recordings were performed intraoperatively under anesthesia—a condition known to alter the latency and morphology of evoked potentials^40^, and the evidence used to derive conclusions was taken mainly from between subjects comparisons.

In parallel, intracranial studies have shown that tonic BHA (50-150 Hz) in posterior perisylvian regions ∼100 ms after stimulus presentation^17^ exhibits the key elements of NCCs: being all-or-none in near-sensory-threshold stimulation, being invariant to report conditions, and showing differences between perceived and non-perceived stimuli in contrastive tasks, as those used for the SAN^21^. Moreover, post-stroke lesions in areas showing this kind of responses produce deficits of tactile awareness^18^.

Given the conceptual, spatial, and temporal similarities between the scalp N140/SAN and the intracranial tonic BHA responses, we hypothesized that they might be two sides of the same coin and tested it with simultaneous high-density EEG and SEEG recordings.

We employed a supra-threshold no-report paradigm for two main reasons. First, to avoid report and task-related confounds. Contrastive near-threshold paradigms with detection reports, though central to the study of conscious perception, may confound awareness-related activity with downstream post-perceptual processes, including decision-making and report^41,42^. Even no-report versions of these paradigms may still recruit cognitive operations linked to task structure, such as attentional control, expectation, working memory, task relevance and saliency^43–46^. Second, in the clinical setting in which these recordings were performed, simultaneous scalp and intracranial acquisition during median nerve stimulation is already technically demanding (Figure 1.a-b), and the inclusion of an active task would have introduced additional equipment requirements, procedural complexity, and recording time. We therefore selected, for this first investigation, the simplest paradigm that both reduced potential confounds and was compatible with the practical constraints of the experimental environment.

BHA is often considered a proxy for neuronal firing, although it may also reflect dendritic processes, and should not be conflated with narrowband rhythmic gamma arising from recurrent excitatory-inhibitory loops^22,23,47^. The time-frequency analysis indeed confirmed that power increases were broadband and mostly located in the 70-140 Hz range (Figure 2.c and Figure 3.i). Our results showed that BHA precedes the LFP by ∼30 ms. This temporal precedence of BHA is in line with the faster nature of the neuronal dynamics that give rise to it, as compared to the spatially and temporally integrated sum of currents that produce the low-frequency LFP^48^. Notably, a recent study has proposed a causal role of high-frequency activity in perceptual decision-making while ascribing a modulatory role to the low-frequency LFP^49^. Crucially, our results also showed that the LFP of contacts displaying tonic BHA is remarkably aligned with the EEG N140/SAN (Figure 2.d), suggesting that the latter is a volume conducted reflection of the former. In addition, the within-subject timing differences and similarities between LFP and EEG were predicted by recording depth (Figure 2.g-h). A plausible biophysical explanation of this effect is that scalp EEG is more sensitive to superficial than to deep generators^50^. Under this view, activity recorded from intracranial contacts located closer to the skull is expected to be represented more directly in the scalp signal, leading to greater similarity and smaller timing differences, whereas signals from deeper contacts are more attenuated and mixed with other sources before reaching the scalp.

In line with the temporal correspondence described above, our results also showed a notable spatial overlap between the whole-brain source-projected EEG activity of 8 subjects with the whole-brain map of sustained BHA responses derived from 99 subjects in a previous study^17^ (Figure 3.b-c). Interestingly, source activity exhibited three volleys with a similar spatial pattern at ∼30, 60, and 120 ms, resembling the BHA response map (Figure 3b) and coinciding, respectively, with the latencies of the phasic, prolonged, and tonic BHA responses reported in that same study^17,21^ (allowing for the BHA-LFP-EEG lag; Figure 2.f). This pattern is consistent with accounts in which recurrent activity is viewed as a necessary^51^, and in some formulations sufficient^52^, condition for conscious perception^27^.

Taken together these results suggest that the N140/SAN is the scalp reflection of the LFP that follows the tonic insulo-opercular BHA response to somatosensory stimulation.

Considering that (i) the tonic BHA in the parietal operculum and insula that we show underlies the N140/SAN during supra-threshold stimulation^17^ also indexes awareness in near-threshold designs^21^, that (ii) its absence is related to perceptual deficits^18^, that (iii) EEG responses to perceived-stimuli in contrastive studies^9,27^ closely resemble the waveforms, latencies, and topographies observed in non-contrastive paradigms, as also seen in our results (Figure 2.a,b,d), and that (iv) responses are not clearly different from baseline in contrastive paradigms when near-threshold stimuli are not perceived (Figure 3.d); our results also suggest that the N140 and the SAN are essentially two experimental manifestations of the same phenomenon. Whether they differ in finer-grained spatial or temporal aspects remains an open question that will require direct within-subject comparisons between paradigms. Such comparisons will need to carefully dissociate the confounds related to the absence of report and possible motor-afference of supra-threshold paradigms; and the motor-, task-, and report-related confounds of contrastive near-threshold paradigms^41–46^; to avoid attributing design-related differences to the NCCs themselves. Notably, it has been shown that the tonic insulo-opercular BHA responses remain invariant regardless of reporting conditions^21^ and that the N140/SAN is modulated by task relevance only with near-threshold stimuli^53^. Most importantly, our results showed a saliency effect that magnified and extended the estimated average responses in frequency and time (Figure 2.c and Figure 3.i-j), even with 5 % of salient stimuli (1/20 trials). This effect did not affect our conclusions (Supplementary Analysis 1); however, experimental paradigms with a higher proportion of salient stimuli than in our case, particularly near-threshold active tasks, may be more susceptible to it^54^. It will be important for future studies to control for this effect and assess how it scales with stimulus intensity. In this context, our findings highlight the importance of establishing synergies between contrastive and non-contrastive approaches for creating better experimental designs.

Once the link between a scalp component and its intracranial generators is established, questions that are otherwise intractable become accessible—either through the high specificity and SNR of direct intracortical recordings or vice versa through the whole-brain coverage of scalp EEG. A central open question regarding the SAN is whether it correlates with conscious perception at the single-trial level^7^. In our data, BHA responses (> 2*σ_baseline_*) to above sensory-threshold stimulations were present in ∼95% of trials while LFP responses in ∼74%. This demonstrates the feasibility of designing experiments to address the question with single trial precision and raises the question of which of these two measures better correlates with perception. If BHA correlates more strongly with perception at the single-trial level, this would suggest that the local activation is the most direct correlate of the process underlying the N140/SAN, with the evoked LFP and EEG reflecting only those instances in which that activity becomes sufficiently synchronized or aligned to generate a measurable field potential. It is also possible that greater baseline variability in the LFP reduces the detectability of otherwise present responses (Figure 1d), thereby necessitating more sensitive methods. The lower single-trial baseline variability of BHA may partly reflect the averaging across frequencies inherent to its computation, which could improve signal stability and, in turn, enhance response detectability.

Bridging across scales also connects their respective bodies of literature. Indeed, our results suggest that the tonic insulo-opercular BHA that distinguishes perceived from unperceived near-threshold stimuli^21^, and its associated LFPs underlie the N140/SAN differences previously reported using scalp EEG^9,10,27^. Likewise, our results showed similar but non-identical saliency effects between BHA and LFP responses, with a faster and a slower approximately two-fold decrease, respectively, which were not evident in scalp data (Figure 3.g). Prior work has proposed hyperbolic decay for the lateralized components of EEG SEPs^29^; here, both BHA and LFP favored an exponential decay. Taken together, these findings illustrate how bridging scales can help characterize brain responses intracranially and extrapolate to the scalp. Moreover, future studies could profit from adding *macro-micr*o^55^ recordings to the simultaneous acquisition setup to also bring the microscopic description into frame (e.g. firing rates), and establish a three-layered hierarchy.

Our analytical framework is readily transferable to the other constituents of the PAN, the Auditory Awareness Negativity (AAN) and Visual Awareness Negativity (VAN), enabling direct tests of shared architecture and precise identification of their sources. More broadly, this approach can be applied to a wide range of tasks and scalp ERPs to resolve generators, compare mechanistic models, and derive intracranial-scalp mappings.

In conclusion, this work helps delineate the intracranial cascade of events giving rise to the scalp N140/SAN and establishes practical and conceptual bridges between the two scales^34^. By linking BHA and LFP dynamics to scalp potentials within individuals, it enables tackling with greater precision questions about localization, timing, and trial-level variability. This cross-scale framework offers a roadmap for future studies to test mechanistic hypotheses about conscious processing, to uncover the mechanisms and generators of other scalp components, and to translate intracranial insights into scalp-level markers for wider clinical and research applications.

## Methods

### Participants

Eight subjects undergoing intracranial monitoring for pre-surgical evaluation of focal drug-resistant epilepsy participated in the study (F = 5; mean age = 33.25; sd age = 7.02). All of them provided written Informed Consent before participating, the study was approved by the local Ethical Committee (protocol number: 463-092018, Niguarda Hospital, Milan, Italy) and it was carried out in accordance with the Declaration of Helsinki. Demographic and clinical information are provided in Supplementary Table 1.

### Median Nerve Stimulation

Stimulation of the median nerve was carried out at the wrist using 1000 constant-current electrical pulses (0.2 ms duration) at 1Hz (Nihon Kohden, Neuropack S3). The stimulation intensity was set as 110% of the motor threshold, which was determined before starting the recording session by increasing the intensity until a thumb twitch was observed (mean intensity = 7.11 mA; sd intensity = 1.59 mA). Stimuli were delivered in blocks of 20 trials (in two sessions blocks comprised 25 trials: S3 and S8; and in one session trials were not divided into blocks: S5). All subjects were stimulated on the side contralateral to the SEEG implant and two subjects were stimulated on both sides, separately (S2 and S4).

### EEG & SEEG Simultaneous Recordings

Simultaneous HD-EEG and SEEG recordings were acquired at the end of the pre-surgical evaluation (Figure 1.a). SEEG signals were recorded at 1000 Hz using a Nihon-Kohden Neurofax-1200 amplifier with DixiMedical (DIXI Medical, Besançon, France) intracerebral electrodes (from 5 to 18 contacts). SEEG electrodes were implanted in specific sites to test one or more neurosurgical hypotheses. For a detailed description of the surgical procedures see^56,57^. HD-EEG signals were recorded at 1000 Hz using an EGI NA-400 amplifier (Electrical Geodesics, Inc; Oregon, USA) with a 256-channel Geodesic Sensor Net (Geodesic Sensor Net; HydroCel CleanLeads). The HD-EEG net was placed on the head by trained neurosurgeons using sterile technique, following a precise step-by-step protocol: (1) sterilization of the net, (2) removal of the protective bandage from the subject’s head, (3) skin disinfection with Betadine and Clorexan, (4) positioning of the HD-EEG net, and (5) reduction of the impedances below 25–50 kOhm using conductive gel, (6) skin disinfection after net removal^58^. The electrical stimulator sent trigger signals recorded by the SEEG and HD-EEG recordings, thus allowing for a precise synchronization of their traces (Figure 1.b).

### Electrode Localization

The positions of the SEEG contacts were determined by registering a post-implant CT (O-arm 1000 system, Medtronic) to a pre-implant MRI (Achieva 1.5 T, Philips Healthcare) by means of the FLIRT software tool^59^ and employing 3D Slicer and SEEG Assistant^60,61^. Normalized coordinates were estimated by performing a non-linear registration between the subject’s skull-stripped MRI and the skull-stripped MNI152 template (ICBM 2009a Nonlinear Symmetric; Figure 1.c) using ANTs’ *SyN* algorithm^62^ and by using Freesurfer’s spherical registration to *fsaverage*. The positions of the HD-EEG electrodes were digitized using a SofTaxicOptic system (EMS s.r.l., Bologna, Italy).

### Data Preprocessing

SEEG and EEG data were imported from the raw Nihon-Kohden and EGI files into Matlab and aligned using the trigger signals delivered by the electrical stimulator employing custom-made scripts. All subsequent pre-processing and analysis steps were performed using custom-made Python scripts based on the *MNE* library^63^: 1) Import, visual inspection, bad channels and segments annotation, and alignment check. 2) SEEG Filtering (high-pass = 0.1 Hz; low-pass = 195 Hz; FIR filter, zero-phase, hamming window, automatic selection of transition band width; Notch filters at 50, 100, 150 and 200 Hz). 3) EEG Filtering (high-pass = 0.5 Hz; low-pass = 40 Hz; FIR filter, zero-phase, hamming window, automatic selection of transition bandwidth). 4) Epochs extraction (from −0.3 to 0.5 s) and baseline correction (−0.1 to −0.025 s). 5) EEG filtering on raw data for ICA (high-pass = 1 Hz, low-pass = 35 Hz; to maximize ICA’s performance). 6) Downsampling to 500 Hz after applying an antialiasing filter at the Nyquist frequency of the new sampling rate. 7) ICA Fitting and Identification (Fastica algorithm with 95% cumulative variance explained for identification of ocular components, i.e. blinks and saccades). 8) Auto-reject^64^, with default parameters, in order to reject epochs and perform single trial and channel interpolation to maximize the amount of retained data. 9) ICA manual rejection. 10) Bad channel interpolation. 11) Final visual inspection and trial rejection. 12) Stimulation artifact removal (using a median filter in a window of 25 ms around 0 and mixing it with the original signal using a Tukey window). 13) EEG average re-referencing. SEEG signals were analyzed in a monopolar scheme with the reference on a silent white matter contact.

### Broadband High-Frequency Activity

Channels showing tonic BHA responses were identified as in a previous study^17^. Briefly, by k-means clustering of gamma-band time courses in responsive contacts (bonferroni corrected t-tests against baseline, successive min = 30 ms, bin length = 10 ms). For each subject, we then selected the channel showing the maximal tonic response within the 50–200 ms window to avoid including neighboring contacts carrying partially redundant information (often 1.5 mm apart; Figure 1.e-f) and to retain a single observation per subject for group level analysis. This choice avoided introducing a nested data structure requiring explicit modeling of within-subject dependence^24^. Given the limited number of channels per subject, we favored this simpler strategy because hierarchical models can be unstable in sparse clustered datasets, would not have materially changed the interpretation of the present results, and are not readily extendable to all analyses used (e.g. cross-correlation).

Single trial SEEG BHA responses were computed by averaging across trials the power extracted using Morlet Wavelets from 50 to 150 Hz with 10 Hz steps and variable number of cycles per frequency (i.e. frequency / 10), z-score normalization with respect to baseline period (from −0.15 to −0.05 s), and averaging across frequencies. In order to compare the topographical patterns obtained with EEG source modeling (see below) with a whole-brain map of BHA, channels showing sustained BHA responses from the previous study with 99 subjects^17^ were plotted on the fsaverage template by taking the distance-weighted sum of responsive contacts in 10 mm spheres (e.g. contacts with distance > 10 mm have weight = 0) and binarizing vertices as with/without responsive contacts using a consensus threshold of 2, to avoid spurious responses of single channels. Of note, the BHA map was created including channels showing either tonic or prolonged responses, which extend to our time window of interest^21^, since EEG is sensitive to both.

### Source Localization

3D models of each subject’s brain anatomy were estimated using Freesurfer^65^. The spatial alignment between the digitized EEG electrodes and the 3D reconstruction of each subject’s head and brain was verified using *MNE*’s coregistration GUI. Source-spaces were created using a recursively divided octahedron (i.e. oct6 spacing). BEM models were created using the following conductivities: scalp = 0.3 S/m, skull = 0.006 S/m, brain = 0.3 S/m. Forward models were created using the aforementioned alignment, source spaces and BEM models. Inverse solutions were computed using the dSPM algorithm with SNR = 5, λ^2^ = 1 / SNR^2^, loose weighting = 1, depth weighting = 1). EEG covariance was calculated in the interval from −0.2 s to −0.05 s using the Shrunk method. Source projections were performed separately for each session, morphed to the fsaverage template, and averaged together. Of note, since SAN/N140 responses are larger contralateral to the stimulated side, sources from sessions in which the right median nerve was stimulated were morphed between hemispheres (*mne.compute_source_morph; xhemi=True*) before being included in the grand average, to align the maximally responsive hemispheres across subjects. The timepoints to plot the source time courses were chosen using *scipy*’s *find_peaks* algorithm on the standard deviation across vertices over time, a metric similar to the GMFP used in sensor-level analysis^25,26^. Each plotted timepoint corresponds to the average ± 10 ms. For comparison with the intracranial BHA map (see above) we employed the top 15 % of reconstructed sources in the window corresponding to the full-width-at-half-maximum of the ROI EEG response (ROI FWHM; 70-150 ms; see below).

### Near-threshold Analysis

The preprocessed epoched data reported in an independent study^9^ was analyzed selecting trials with intensity values 5 and 6 (i.e. near-threshold) from the matching task (MT). Signals were re-referenced to the average and condition-wise evoked responses were computed for undetected (det = 0) and detected (det = 1) trials for each subject. A ROI trace was obtained from each subject using channel C6 as in the original article. Statistical comparisons were performed using cluster-based one-sample permutation tests across time with 5000 permutations. The analyses tested the *detected* - *undetected* (det1 - det0) contrast as well as each condition against baseline, with a cluster-forming threshold of *p* < 0.05, to maximize sensitivity. Baseline correction before testing employed the whole available pre-stimulus interval (from −51 to −1 ms).

### Statistical Analyses

The ROI for EEG analyses were taken from previous literature^9,10,27^. The temporal window of interest, for analyses and images that required it, was computed by finding the peak of the average EEG response in the ROI using *scipy*’s *find_peaks* algorithm and calculating the Full Width at Half Maximum (ROI FWHM).

To assess the temporal relationship between BHA and LFP and between LFP and EEG, we smoothed the single session signal using a moving average of 30 ms and extracted the peak latency (latency of maximum for BHA, and latency of minimum for EEG and LFP, due to the negative polarity). In the two subjects that received stimulation to both the right and left median nerve values were averaged to avoid violating the assumption of independence of observations of the subsequent statistical tests. Both comparisons were performed using paired t-tests with Bonferroni correction for multiple comparisons. To test the relationship between timing variability and proximity we employed a quadratic regression, since temporal differences could be either positive or negative. We also verified that results remained unchanged when using a linear regression on absolute difference values (data not shown). To assess the relationship between the LFP-EEG similarity and proximity we first computed the cross-correlation (ρ) between the signals (80 to 250 ms, maximum lag = 60 ms) at the single session level and extracted the maximum correlation and the corresponding lag^66^. Values were then averaged by subject and tested with a linear regression analysis.

Similarity between the average reconstructed sources (70-150 ms; ROI FWHM) and the intracranial BHA map was assessed using the Dice-Sørensen coefficient (i.e. Dice Similarity Index), a common metric of similarity employed in medical image analysis^67^. Statistical significance of the similarity was assessed by creating a null distribution using 5000 spatial permutations (i.e. random spherical rotations that preserve autocorrelation)^68^ employing *neuromaps*^69^, and computing *p-values*^70^ as (*count*{*DiceNull* > *Dice*} + 1) / (*Nperm* + 1).

Single trials were considered as responsive if the peak of the signal from 50 to 200 ms exceeded 2 times the standard deviation of the baseline (−150 to −50 ms). The comparison in the proportion of responsive trials between BHA and LFP was performed using a Wilcoxon signed-rank test, to avoid the normality assumptions of parametric tests.

Saliency/habituation was assessed by computing the average by trial position in the interval from 70 to 150 ms (ROI FWHM). Data from the session without block design was not used for this analysis and data from the two sessions with blocks of 25 trials were considered up to the 20^th^ trial of each block. Grand averages by trial position were obtained by combining subject-level data through a weighted average according to the number of trials used for each subject and position. This analysis was performed at the group level since with the number available trials EEG data lacks SNR to perform accurate within-subject across-trials quantifications. Data was expressed as percentage with respect to trial in position one. Two models were fitted to each data type (BHA, LFP and EEG), an exponential *f*(*x*) = *a* + *b* · *exp*(− *c* · *x*)) and a hyperbolic decay function (*f*(*x*) = *a* + *b*/*x*), using *scipy*’s curve fitting tools, following previous reports^29^. Fits were evaluated through adjusted R^2^, ΔAICc (small sample size modification of Akaike Information Criterion)^71^, Wald *p-values* from t-statistics, visual inspection of Q-Q plots of standardized residuals and Shapiro-Wilk tests for normality (Supplementary Figure 4 and Supplementary Table 7).

All analyses reported above were repeated but only considering trials 2-20 (TP-20) to verify that results remained unchanged when excluding the salient first trial of each stimulation block (Supplementary Analysis 1).

Time–frequency, BHA, LFP, and EEG responses were compared between the subject-averages of trials in all positions (TP 1-20) and trials of trials in positions 2-20 (TP 2-20) using non-parametric cluster-based one-sample permutation tests with 5000 permutations. For the time–frequency data, condition differences were tested across the full time–frequency plane using a spectro-temporal adjacency matrix, whereas the BHA, LFP, and EEG time courses were tested across time. In all analyses, clusters were defined with a two-tailed cluster-forming threshold of *p* < 0.01. The analysis window ranged from −100 ms to 400 ms relative to event onset to avoid the edge artifacts of the time-frequency transform.

## Data and Code Availability

The dataset and code required to reproduce the results of the study are publicly available at:

[link; see NOTE].

Data for the near-threshold analysis was obtained from an independent study^9^ and is also publicly available (see above):

https://doi.org/10.6084/m9.figshare.13160381

NOTE: doi and link to be added upon acceptance of the article.

## Acknowledgements

This research was funded by the Progetto Di Ricerca Di Rilevante Interesse Nazionale (PRIN, P2022FMK77), Ministry of University and Research (MUR); by the Italian Ministry of Health—(Ricerca Corrente 2022–2024); by the Department of Philosophy “P. Martinetti” of the University of Milan under the Project “Departments of Excellence 2023-2027” awarded by the Ministry of University and Research (MUR); by the Italian National Recovery and Resilience Plan (NRRP), M4C2, funded by the European Union - NextGenerationEU (Project IR0000011, CUP B51E22000150006, “EBRAINS-Italy”); by the European Research Council HORIZON-INFRA-2022 SERV (Grant No.101147319), “EBRAINS 2.0: A Research Infrastructure to Advance Neuroscience and Brain Health”; and by the European Research Council (ERC-2022-SYG - 101071900 - NEMESIS).

## Author contributions

E.M.: conceptualization, design, analysis, interpretation, software, data management, draft and revision; F.M.Z.: patient management, acquisition and revision; S.R.: interpretation, draft, revision; F.M.: software and data management; M.D.V.: interpretation, draft and revision; D.A.: acquisition, interpretation, draft and revision; P.dO.: clinical supervision and patient management; L.N.: conceptualization, clinical supervision and patient management; I.S.: clinical supervision, patient management, acquisition and revision; M.M.: conceptualization, design, interpretation and revision; P.A.: conceptualization, design and interpretation; A.P.: conceptualization, design, interpretation, draft and revision.

